# Deep Sequencing revealed ‘Plant Like Transcripts’ in mosquito *Anopheles culicifacies*: an Evolutionary Puzzle

**DOI:** 10.1101/010009

**Authors:** Punita Sharma, Swati Sharma, Ashwani Kumar Mishra, Tina Thomas, Tanwee Das De, Sonia Verma, Vandana Kumari, Suman Lata Rohilla, Namita Singh, Kailash C Pandey, Rajnikant Dixit

## Abstract

As adult female mosquito’s salivary gland facilitate blood meal uptake and pathogen transmission e.g. *Plasmodium*, virus etc., a plethora of research has been focused to understand the mosquito-vertebrate-pathogen interactions. Despite the fact that mosquito spends longer time over nectar sugar source, the fundamental question ‘how adult female salivary gland’ manages molecular and functional relationship during sugar vs. blood meal uptake remains unanswered. Currently, we are trying to understand these molecular relationships under dual feeding conditions in the salivary glands of the mosquito *Anopheles culicifacies*. During functional annotation of salivary transcriptome database, unexpectedly we discovered a cluster of salivary transcripts encoding plant like proteins. Our multiple experimental validations confirmed that Plant like transcripts (PLTs) are of mosquito origin and may encode functional proteins. A comprehensive molecular analysis of the PLTs and ongoing metagenomic analysis of salivary microbiome provide first evidence that how mosquito may have been benefited from its association with plant host and microbes. Future understanding of the underlying mechanism of the feeding associated molecular responses may provide new opportunity to control vector borne diseases.

## Introduction

Sugar feeding by adult mosquitoes is essential for regular metabolic energy production to maintain a wealth of behavioral, structural, and physiological demands. While blood feeding by adult female mosquitoes is essential to meet the extra nutrient requirement for egg production and life cycle maintenance. Thus blood and sugar feeding are mutually exclusive and antagonistic behavioral-cum-physiological properties of the conflicting demands(Foster, 1995). It is worth of interest to understand the biological consequences of the mosquito tissues e.g. salivary glands, midgut etc. involved in feeding and digestion.

Adult female mosquito salivary glands initiate first biochemical communication to facilitate sugar as well as blood feeding. Additionally, it also potentiates pathogen transmissions and therefore research has largely been focused to understand the role of salivary glands in relation to blood feeding(Das et al., 2010; Dixit et al., 2009; James, 2003; Ribeiro et al., 2010; Rodriguez and Hernandez-Hernandez Fde, 2004). Despite the mosquitoes spend longer time over plant (floral nectar) sugar source, several fundamental questions in relation to the evolution of the dual feeding behavior ‘in general’ and functional relationship of salivary glands during sugar vs. blood meal uptake ‘in specific’, remain unanswered. It is believed that dual feeding behavior evolution might have occurred from herbivores feeding behavior to blood acquisition (Lehane, 2005)

Currently we are investigating the salivary associated molecular factors that affect the mosquito feeding behavior and *Plasmodium* transmission(Dixit et al., 2011; Dixit et al., 2009). During ongoing annotation of our recent salivary transcriptome database (unpublished) of the Indian malarial vector *Anopheles culicifacies*, we unexpectedly observed a cluster of Plant like transcripts (PLTs) in the sugar fed library. A comprehensive molecular analysis of the PLTs and ongoing metagenomic analysis of salivary microbiome provide initial evidence that how mosquito evolved and adapted for feeding over plant host.

## Results & Discussion

In recent years, next-generation sequencing has not only opened the door for functional genomics analysis, but also emerging as an important tool to understand the evolutionary relationship of the molecular codes identified from non-model organisms(Bao et al., 2012; Gibbons et al., 2009; Hittinger et al., 2010; Su et al., 2012; Wang et al., 2010). Accordingly, we adopted Illumina based deep sequencing approach as a proof of concept for gene discovery tool. We sequenced two cDNA libraries prepared from the salivary glands, collected from 3-4 days old either sugar or blood fed (within 1hr of blood feeding) adult female mosquitoes. This protocol in fact generated a total of ∼58.5 million raw reads, which were quality filtered and *denovo* assembled, yielding a set of 11,498 (5808 for sugar fed (SF) & 5690 for blood fed (BF) library) contigs. Initially, the quality of the assembly was carefully examined by multiple homology search analysis of the whole transcriptome dataset against draft genome/transcript databases for the mosquito *A. culicifacies*, available at www.vectorbase.org.

As expected ninety two percent transcripts yielded significant match (10^−5^ e-value) to the draft genome of the mosquito *A. culicifacies*, at nucleotide level. Later, we selected few full length cDNA transcripts (>1000bp) and compared them with previously well annotated genes identified from other mosquitoes (S1). Subsequent validation of the selected transcripts by RT-PCR based expression analysis not only confirmed the quality of the assembly, but also allowed us to find out those rare Plants like transcripts which remained previously un-noticed, as mentioned below. Detail stats of the salivary transcriptome assembly kinetics have been summarized in the **S1.**

### Pilot discovery of Plant like transcripts

BLAST2GO analysis of both the libraries independently showed distinct annotation kinetics (S2). Unexpectedly, species distribution analysis revealed several ‘Plant Like Transcripts’ (472 PLTs/∼8%), with highest similarities (>95% identity) to the ‘plants species’ in the sugar fed, but absent in the blood fed transcriptome database **(Fig.1a/ST-1)**. To find out the possible evolutionary relationship of putative PLTs, we performed extensive BLASTX analysis against either NR database or Insect specific database at NCBI. From this analysis we further characterized three categories of transcript(s) (i) one transcript: encoding highly conserved alpha-tubulin (cytoskeleton associated protein), showing highest identity (>95%) to plant and (85-90%) identity to insect; (ii) two transcripts: encoding aquaporin (water channel membrane protein) (Maurel et al., 2008) and active site of the cysteine protease (protein chewing enzyme)(Grudkowska and Zagdanska, 2004) showing highest (>90%) identity to plant and 40-52% identity to insect (iii) two transcripts: encoding dehydrin (cold stress response protein) (Hanin et al., 2011) and expansin (plant cell wall loosening protein)(McQueen-Mason and Cosgrove, 1995) only matched to plants, but remained unmatched to any insect database **(S3a,b,c).**

**Fig. 1:**
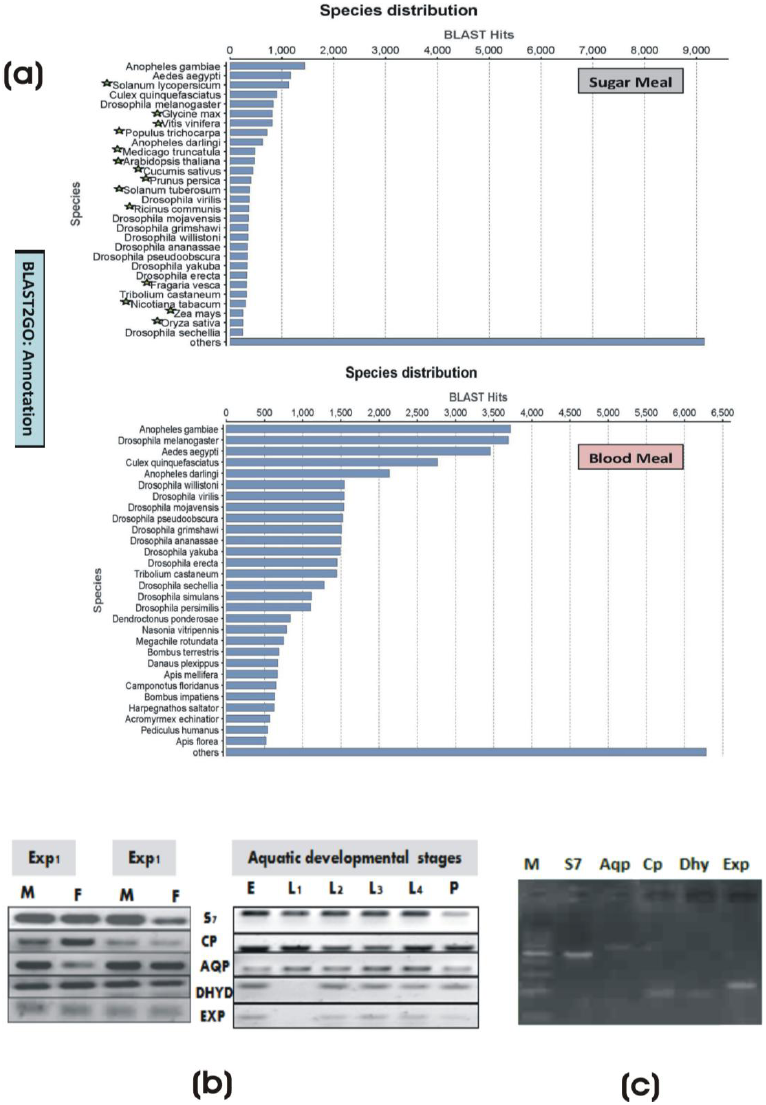
Mosquito encodes Plant like proteins: (a) BLAST2GO based Species distribution analysis in response to sugar and blood feeding. Green star mark indicates the name of Plant species, best match to the NR database in the sugar fed, but absent in the blood fed salivary transcriptome. (b) Confirmation of the nature of Origin: RT-PCR expression of PLTs during aquatic development of the mosquitoes.

The above fascinating observations prompted to know whether such PLTs have previously been predicted or identified from any other mosquito species. To clarify, we performed extensive homology search and surprisingly find out one ESTs dataset of *Aedes aegypti* larval cDNA library, where several PLTs have been recorded and submitted in the unigene database at NCBI (http://www.ncbi.nlm.nih.gov/unigene?term=Aae.16606&cmd=DetailsSearch/S4a/b). We believe such putative PLTs remain undescribed, probably due to tissue contamination suspect (Venancio et al., 2009). Nevertheless, in our case we collected the salivary tissues from the same cohort of the mosquitoes offered sugar or blood meal, and processed both the samples for library sequencing in identical conditions. Thus, we believe the absence of PLTs in blood fed transcriptome; nullify the chances of any suspect of contamination. However, above observations still rose several puzzling, but important arguable questions: (i) do these PLTs really express in mosquito tissue (ii) if express, do these transcripts have any evolutionary significance in relation to feeding preference and adaptation?

## PLTs are of mosquito origin

First, we did a deep enquiry with technical staff and confirmed that under standard rearing facilities, mosquitoes are never exposed to any plant material. However to rule out the possibilities of any contamination, for each experimental analysis, we separately maintained the experimental mosquitoes as detailed in the methodology section. For technical validation of the PLTs origin, we conducted a series of experiments: (i) in two independent experiments, first we examined and verified the RT-PCR based expression of at least 10 selected PLTs (Fig1b/S5a), in the sugar fed salivary glands of adult male and female mosquitoes; (ii) interestingly, we also observed that PLTs expression is not only restricted to the mosquito tissues, but also express during the aquatic developmental stages viz. egg, larva, and pupa of the laboratory reared mosquitoes **(Fig.1b).** Here, it is also important to mention that mosquito egg and pupa stages are metabolically active, but never feed; (iii) we also observed similar amplification of selected PLTs through genomic DNA PCR (Fig.1c); (iv) we further carried out the functional validation of one of the plant homolog PLT encoding dehyrin protein, by Real-Time PCR as well as Immunoblot analysis (see fig. 4a/b); (v) lastly, from ongoing annotation of another independent transcriptome sequence database originated from non-salivary tissue of adult female mosquito *Anopheles culicifacies* (unpublished), we again observed similar PLTs (see S5b). Taken, together our experimental data strongly validate that mosquito genome may code plant like proteins. The poor match of PLTs to the available *A. culicifacies* mosquito genome sequences, could be due to following possible reasons: (i) strain specificity i.e. we worked on Indian strain, while available sequence data are originated from wild caught Iranian strain of *A. culicifacies* (www.vectorbase.org; Neafsey et al., 2013); (ii) incomplete annotation i.e. the quality and annotation of draft genome of Iranian strain may be incomplete (iii) genome assembly pipelines i.e. it is usual practice to filter out poorly matched, non-linage specific orphan genes etc. as ‘junk DNA’, during genome assembly from raw data; (iv) we believe that observed PLTs in our transcriptome screen may be an added benefit of deep sequencing approach which allowed to recover the rare transcripts, but may not be easy to recover in case of genome sequence.

It has long been accepted and proved that a significant variation exists in the chromosomal DNA as well as genome size within *Anopheline* and other mosquito species(Rai, 1999; Neafsey et al., 2013), but how these variations differentially affect the mosquito biology viz. behavior, physiology, immunity and vectorial capacity etc., are poorly understood at molecular level. Thus, we believe that unexpected finding of PLTs may be one of the valuable source databases to improve our knowledge and understanding the biological meaning of complex genomic variations within mosquito species.

### Phylogenomic analysis of Plant like transcripts

Excitingly, the above data confirmation, prompted to follow up the associated evolutionary consensus, favoring plant-mosquito relationship: a parallelism setting where different species from unrelated taxa faces the common selective pressure (Zhen et al., 2012). Initial multiple sequence alignment analysis revealed significant heterogeneity (substitution/deletion) of amino acid residues, but also indicated unique conservation of insect or plant specific residues within the mosquito *A. culicifacies,* result a clade formation with plant species **(Fig.2a/b/S6)**. Subsequently, we also tested whether evolution of common traits from unrelated taxa owing to similar selection pressure favors adaptive significance.

A maximum likelihood (ML) estimation was applied to calculate and compare the sitewise likelihood (ΔSSLS) values between two hypothesis i.e. mosquito-mosquito species evolution (H_0_) and mosquito-plant convergent adaptive evolution (H_1_), for the selected PLTs. The site wise log likelihood plot indicator i.e. divergence towards negative (ΔSSLS) was compared with LRT (likelihood ratio test), using parametric bootstrap at 1000 replicate analysis (cut off p-value 5%). Final data analysis and comparison stats favored the convergent hypothesis, (Arendt and Reznick, 2008) demonstrating that mosquito *A. culicifacies* PLTs followed convergent model favoring (H1), an adaptive evolution for sugar feeding associated functional relationship with plants **(Fig.2c/S6).** Our analysis also support the previous observations noted for the evolution of echolocating gene clusters among bats and bottlenose dolphins(Parker et al., 2013). Additionally, the predicted 3D structural analysis revealed fine conservation of the active functional domains in the mosquito and plants proteins e.g. cysteine protease **(Fig.2d/S7)**. From these studies, we concluded that mosquito feeding associated genes are not only evolving actively, but also acquiring new genes (e.g. dehydrin, expansin), to adapt successfully over plant host.

**Fig. 2:**
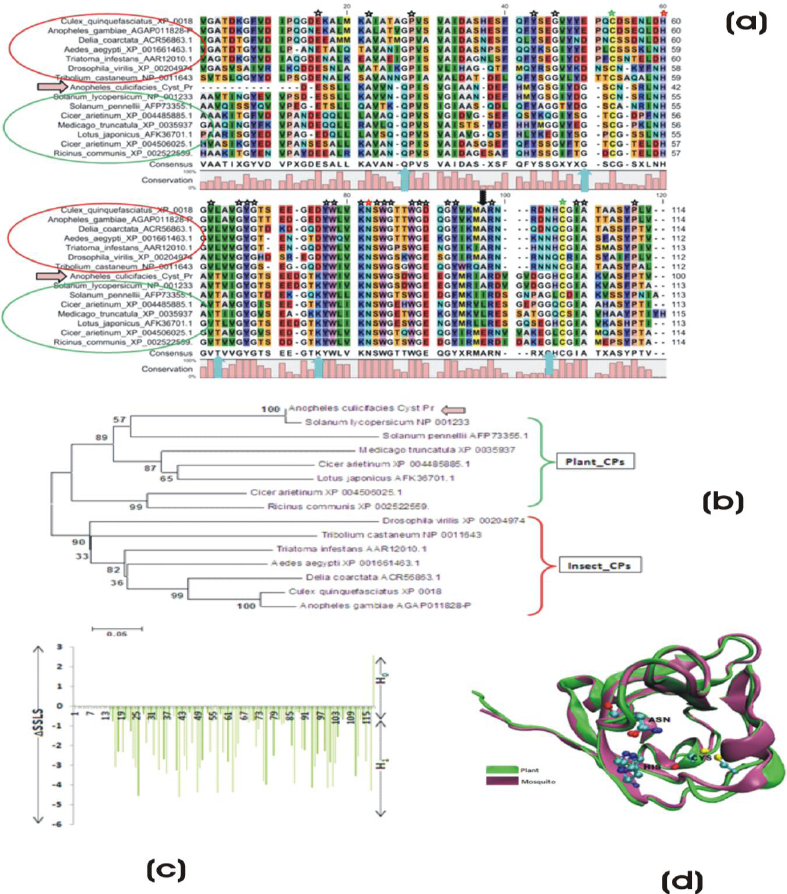
Feeding associated molecular complexity of the mosquito salivary transcripts: **(a)** Molecular analysis of partial cDNA sequence encoding (100AA) Plant-like Cysteine protease active domain: Multiple sequence alignment showing molecular relationship of AcSgCp with plant (Green circle) as well as insects (Red circle) cysteine proteases: conserved residues (marked as 
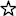
) as well as conserved active site residue (marked as 
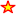
). Green 
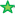
represents conserved cysteine residues, which enables disulfide formation. Upward arrow mark 
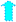
 represent unique plant specific amino-acids residues also conserved in the *Anopheles culicifacies,* while downward arrow mark 
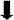
 represent unique insect specific residue conserved in *A. culicifacies* and only in *Solanum Lycopersicum*. **(b)** The evolutionary history of AcSgCp inferred using the Neighbor-Joining method, favoring a clade formation with *S. lycopersicum* and other plant cysteine proteases. **(c)** Relationship between strength of convergent evolution favoring adaptive significance of feeding associated PLTs: A maximum likelihood (ML) estimation was applied to calculate and compare the sitewise likelihood (ΔSSLS) values between two, species evolution (H0) and convergent adaptive evolution (H1) hypothesis, for Cysteine protease (see text for details). **(d)** Structural comparison between predicted 3D structure of the mosquito, and solved structure of the plants cysteine protease: Aspargine (ASN) and Histidine (HIS) indicate conserved residue of the active site.

#### Feeding associated molecular complexity of ‘salivary-sugar-microbe’: A tripartite interaction

Evolution of herbivores insect-plant association represents one of the dominant interactions over millions of years(Agrawal et al., 2012; Ehrlich, 1964; Fraenkel, 1959). These interactions are thought to play an important role in the co-evolution of molecular effectors arms, enabling effective adaptation over each other (Hogenhout and Bos, 2011). Uncovering of the molecular mechanisms of the herbivores insect-plant interaction has greatly facilitated the design of molecular strategies to save valuable crops from insect pests (Baldwin et al., 2001; Felton and Tumlinson, 2008; Ferry et al., 2004; Maffei et al., 2007). However, such studies have not been given special attention to mosquitoes. From the unexpected findings of the mosquito PLTs, we interpreted that either studies in relation to the sugar feeding associated salivary biology has largely been ignored (Dixit et al., 2011; Juhn et al., 2011) or the mosquito *A. culicifacies* may have evolved with more complex genetic architecture favoring evolution of environmentally-guided several traits viz. carbon metabolism; light mediated photo conditions for mating, feeding, survival etc. Therefore, to predict sugar metabolism associated molecular and functional relationship of salivary PLTs, initially we analyzed all the putative plant like transcripts against three databases (Reactome; KEGG and Biocycles) annotated for *Arabidopsis thaliana*, using KOBAS online software (http://kobas.cbi.pku.edu.cn/home.do).

Notably, we observed that 18 salivary transcripts encoding proteins related to at least five Biocyclic pathways linked to photosynthetic organelles viz. plastid in plants **(S8a/T-1; see also S2)**. To verify above predicted ‘plastid’ related salivary transcripts, Fisher’s exact test analysis identified a pool of 11 transcripts differentially expressed in the sugar fed mosquitoes (Fisher test p<0.001; **S8b.**); encoding important enzymes/proteins, associated with one of the key pathway “Carbon fixation in Photosynthetic Organisms” **(Fig.3a)**. Further, we also identified four unique salivary transcripts encoding different enzymes linked to three other secondary metabolite synthesis pathways:-namely ‘Trepenoid Backbone Biosynthesis’ (4-hydroxy-3-methylbut-2-enyl diphosphate reductase/E.C.1.17.1.2, LYTB); ‘Carotenoid Biosynthesis’ (Phytoene Synthase/E.C.2.5.1.32, PS); and ‘Flavonoid Biosynthesis’ (3-dioxigenase/E.C.1.14.11.9 & 3’ beta-hydroxylase/E.C.1.14.13.88) pathways restricted to the plants (S9). A comprehensive molecular and phylogenetic analysis of few selected transcript, encoding an enzyme 4-hydroxy-3-methylbut-2-enyl diphosphate reductase/E.C.1.17.1.2 (LYTB) and phytoene synthase/E.C.2.5.1.32 (PS); exclusively revealed unique evolutionary relationship to the cyanobacteria, algae, plants and aphid *Acyrithosiphon pisum* **(Fig. 3b,c S9)**.

**Fig. 3:**
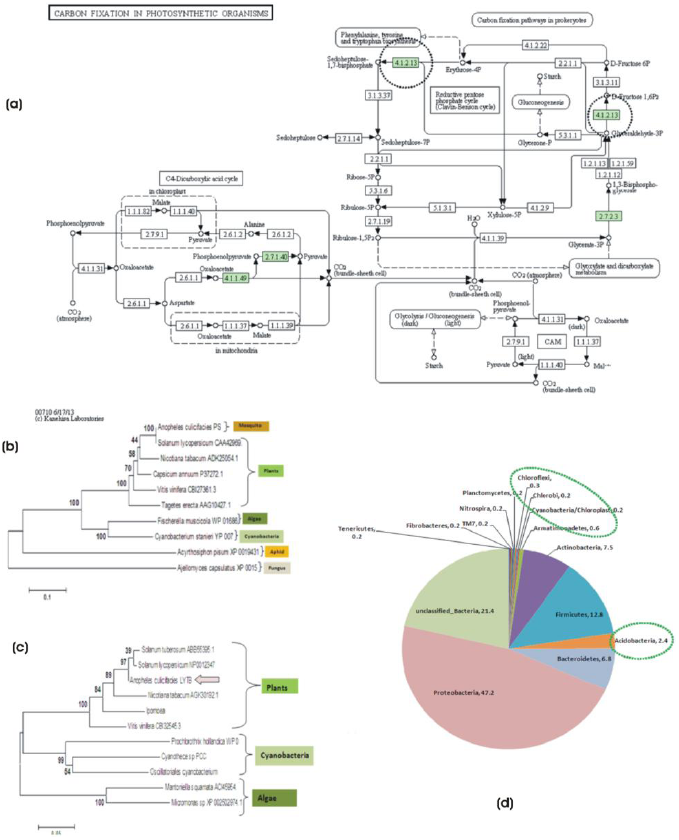
Molecular Evidence that mosquito encodes plant like photosynthetic machinery components partly shared by symbiotically associated salivary bacteria for carbon fixation and metabolism: **(a)** KEGG prediction of salivary transcripts (differentially expressed/ Fisher test p<0.001) encoding enzymes (Green) involved in “Carbon Fixation in Photosynthetic Organisms” pathway known to be restricted to the photosynthetic plants organelles e.g. plastids only (see text). **(b)** Phylogenetic analysis of a unique mosquito salivary transcript, encoding a Plant homolog 4-hydroxy-3-methylbut-2-enyl diphosphate reductase/E.C.1.17.1.2 linked to the “Trepenoid Backbone Biosynthesis” pathway. **(c)** Phylogenetic analysis of a unique mosquito salivary transcript, encoding a Plant homolog Phytoene Synthase/E.C.2.5.1.32 linked to the “Carotenoid Biosynthesis” pathway. In fact like other animals, insects are also believed to absorb carotenoid pigment (an eye pigment) from plant food. Additionally, lower microbes such as algae and cyanobacteria also carries LYTB/PS gene in their genome. Phylogenetic analysis of the salivary LYTB & PS showed unique association with the plant, as well as microbial LYTB while PS also showed evolutionary relationship to the novel PS gene recently identified from sap sucking insect *Acyrithosiphon pisum*, suggesting that mosquito LYTB/PS might have evolved, for independent synthesis of the carotenoid synthesis assisting feeding adaption preference over plant host. **(d)** Identification of symbiotically associated salivary microbial flora predominated and unique bacteria (marked green circle), probably assisting mosquito to adapt, feed and metabolize diverse carbon rich sugar sources of plant origin (see another report for detail).

In fact during its early development mosquito larvae start to feed diverse micronutrients e.g. bacteria, algae, fungi etc., and switch to feed on nectar sugars in adult mosquito stage. Thus, it could be possible that a longer association and regular microbes-mosquito-plant interactions(Bennett, 2013; Pieterse and Dicke, 2007), might have favored insect/mosquito to adapt, feed, digest sugar and selective synthesis of secondary metabolites/pigments, essential for specific phenotype e.g. visual pigmentation/dark body coloration(Benedict and Seawright, 1987). Our recent metagenomic analysis of salivary microbiome identified several diverse unique bacterium phyla including Chlorobium, Cyanobacteria, Nitrospira and other phototrophic bacteria associated with salivary glands **(Fig.3d)**, but absent in the gut of the laboratory reared 3-4 days old adult female mosquito *Anopheles culicifacies* (Sharma et al., 2014).

These findings further support the hypothesis that mosquito may have feeding associated distinct plant like molecular machinery components, partly shared by residing symbiotic bacterial community for diverse carbon/nitrogen rich plant sugar source metabolism. For example, finding of prominent salivary associated Acidobacteria (2.4%), may facilitate the utilization of plant polymer viz. cellulose/xylan sugars of diverse origin(Eichorst et al., 2011), as reported in the gut of the wood feeding larvae of Huhu Beetle (*Prionoplus reticulari*)(Reid et al., 2011). Furthermore, recent study on light-induced ATP synthesis from the chloroplastidic-like carotene pigments in *’Acyrithosiphon pisum*’, a plant sap sucking aphid, provides first molecular evidence that aphid genome may carry plant like photosynthesis machinery components(Valmalette et al., 2012). A fungal mediated lateral horizontal gene transfer mechanism has been proposed for the evolution of carotenoid biosynthesis gene in this aphid(Moran and Jarvik, 2010).

Although, accumulating evidences of genetic material transfer within metazoan are still at premature stage but strongly suggest that acquisition of new beneficial traits, may favor improved survival and adaptation values in changing ecologies (Boto, 2014). Thus we believe our finding may begin to unlock previously unexplored biology, to rebuild new hypothesis “how insects are most successful” to feed, adapt and survive in the diverse environments. In support of these observation, next we attempted to validate whether PLTs encodes a functional protein for specific function (see below).

### Mosquito encoded Plant-homolog Dehydrin: a functional validation

Dehydrins are low-temperature (LT) acclimation and evolutionarily conserved proteins that allows to develop efficient tolerance to drought and cold stress among photosynthetic as well as in some non-photosynthetic organisms such as yeast (Campbell and Close, 1997; Close and Lammers, 1993; Li et al., 1998; Mtwisha et al., 1998). Dehydrins are characterized by conserved K-segment comprising consensus KIKEKLPG amino acid sequence towards the C-terminus and may be repeated one to many times to encode 9 -200 kDa protein (Close, 1997; Close et al., 1993; Ouellet et al., 1993; Takahashi et al., 1994).

Unlike plants, insect dehyrin have not been reported so far, though a putative transcript AGAP000328 has been predicted from mosquito *A. gambiae* genome, carrying (PF00257 domain) a signature of dehyrin like proteins **(S10)**. In our RT-PCR analysis, we observed a constitutive expression of *AcDehydrin*, throughout the aquatic developmental stages of the mosquito, indicating that identified PLT *AcDehydrin* transcript may encode a putative functional protein. Thus we characterized mosquito encoded plant homolog protein dehydrin in detail **(S10)**. Our real-time PCR analysis repeatedly confirmed that dehydrins highly express in the egg than larva or pupae **(Fig. 4a/S10),** suggesting an abundant accumulation in the egg.

**Fig. 4:**
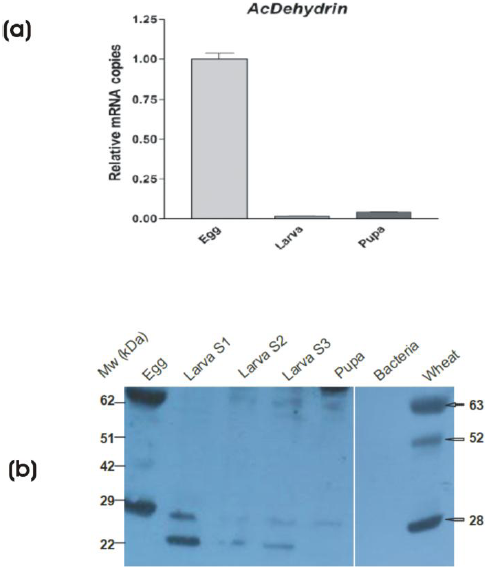
Functional validation of mosquito encoded Plant homolog Dehydrin: **(a)** Real-Time PCR analysis of Dehydrin, demonstrating abundant expression in the mosquito egg, **(b)** Immunoblot analysis of Dehydrin expression during the development of the mosquito: Anti-dehydrin antibody recognize three protein bands of expected size (28,52 & 63kDa) in the control wheat seedling samples (arrow mark). Mosquito samples included Egg, Larval stages (S1,S2,S3) and pupa while negative reference includes bacterial protein sample.

For functional validation of *AcDehydrin* protein, we examined the developmental expression of the dehydrin protein through immuno-blotting assay using anti-dehyrin antibody (kind gift from Dr. Timothy Close). In these experiments we used wheat seedling protein sample as positive reference control, while bacterial protein sample as negative reference. The anti-dehydrin antibody not only recognized the expected (28, 53 and 62 kDa) protein band in the wheat samples(Borovskii et al., 2002), but also identified two proteins (28 and 62kDa) abundantly expressing in the egg as compared to other developmental stages of the mosquito **(Fig.4b)**. Taken together, our functional validation suggests that mosquito encoded *AcDehydrin* protein may have functional role similar to plant.

Like other late embryogenesis abundant proteins (LEA), dehydrins accumulate to high amounts in plant embryos, but remains undetectable in other vegetative tissues until their exposure to cellular dehydration stress. The stress exposure results in their rapid induction and binding to multiple proteins, probably through intramolecular hydrogen bonding to protect tissue damage from dehydration/cold stress (Hanin et al., 2011).We hypothesize that mosquito *A. culicifacies*, may have survival benefits of cold stress tolerance. Future studies involving dsRNA mediated gene silencing approach may unravel molecular and functional relationship of the PLTs, controlling feeding and adaptation phenotypes in the mosquito (Lu et al., 2012; Scott et al., 2013).

## Material & Methods

**Mosquito Rearing:** Cyclic colony of the mosquito *Anopheles culicifacies* sibling species A, were reared and maintained at 28 ±2°C /RH 80% in the insectary fitted with a simulated dawn and dusk machine, essentially required for proper mating and feeding at NIMR (Adak et al., 1999). All protocols for rearing, maintenance of the mosquito culture were approved by ethical committee of the institute. For our specific research work, we harvested 80-100 pupa and allowed them to emerge in standard mosquito cages. For regular sugar supply the mosquitoes were offered sterile 5% sugar solution (crystal sugar dissolved 5g/100ml water) using cotton swab, while blood meal was offered to adult mosquitoes through rabbit. For aquatic development, gravid female were allowed to lay egg on moistened filter paper mounted inside small plastic cup (e.g. ice cream cup), semi-filled with water. Hatched larvae were feed on mixed dried powder of fish food and yeast.

**cDNA library preparation & sequencing:** Total RNA isolation and double stranded cDNA library preparation was done by PCR-based protocol as described previously(Dixit et al., 2009). Briefly, DNase-treated total RNA was reverse-transcribed and amplified to make double-stranded cDNA (ds-cDNA) through a PCR-based protocol using SMART cDNA synthesis kit (catalogue No. 634902; BD Clontech, Palo Alto, CA, USA), following the manufacturer’s instructions. For deep sequencing, Single-End RNA-seq protocol was used for each sugar fed and blood fed salivary gland tissue libraries, by commercial service providers (NxGenBio Life Sciences, New Delhi, India). The tagged Single-End RNA-Seq libraries were diluted and pooled in equimolar concentrations and sequenced using TruSeq^TM^ SBS Kit V2 on Illumina GAIIx (Illumina, San Diego, CA) for generating 1x36bp single end sequencing reads.

**Transcriptome assembly:** Following sequencing, the low quality bases were filtered or trimmed using in-house Perl scripts. All the bases, above Q20 phred score were used for further downstream analysis. De-novo transcriptome assembly was performed using Trinity assembler (Grabherr, 2013) with the default settings *k*-mer size of 25, minimum contig length of 200, paired fragment length of 500, 16 CPUs, with butterfly Heap space of 100G (allocated Memory). Prior to submission of the data to the Transcriptome Shotgun Assembly Sequence Database (TSA), assembled transcripts were blasted to NCBI’s UniVec database (http://www.ncbi.nlm.nih.gov/VecScreen/UniVec.html) to identify segments with adapter contamination and trimmed when significant hits were found. This adapter contamination may result from sequencing into the 3' ligated adapter of small fragments (<100 bp).

**Functional annotation:** Following *DENOVO* clustering, CAP3 assembly using desktop cDNA annotation system (Guo et al., 2009) was used to build final conting/transcripts dataset for functional annotation. The assembled transcripts were subjected to similarity search against NCBI’s NR database using the BLASTx algorithm (Altschul et al., 1990), with a cut-off E-value of ≤10^-3^ using BLOSUM62 matrix as well as GO annotation/Interproscan analysis using BLAS2GO(Conesa et al., 2005). Biocyclic pathway analysis for PLTs KOBAS online (http://kobas.cbi.pku.edu.cn/home.do) software (Xie et al., 2011).

**PCR based gene expression analysis:**The desired tissues viz. salivary glands, midgut and hemocyte (Rodrigues et al., 2010) or the whole body, were directly in the Trizol. Total RNA was isolated using standard Trizol method, followed by first-strand cDNA synthesis using Oligo-dT or Random Hexamer primers (Verso kit). For differential expression analysis, routine RT-PCR and agarose gel electrophoresis protocols were used. Relative gene expression was assessed by QuantiMix SYBR green dye (Biotool Biolabs, Madriad, Spain) in Eco-Real-Time PCR Machine (Illumina). PCR cycle parameters involved an initial denaturation at 95°C for 15 min, 40 cycles of 10 s at 94°C, 20 s at 55°C, and 30 s at 72°C. Fluorescence readings were taken at 72°C after each cycle. A final extension at 72°C for 5 min was completed before deriving a melting curve, to confirm the identity of the PCR product. Actin gene was used as an internal control in all qPCR measurements, where minimum two technical replicates were used in each Real-Time experiment. To better evaluate the relative expression, each experiment was performed in three independent biological replicates. The relative quantification results were normalized with internal control Actin gene and analyzed by 2^−ΔΔCt^ method (Livak and Schmittgen, 2001).

**Phylogenomics analysis:** Following primary BLASTX analysis, the reference sequences from the selected top hits were retrieved and edited for subsequent analysis in the FASTA format. Multiple sequence alignment was performed using ClustaX2 (Larkin et al., 2007). The CLC Sequence viewer (http://www.clcbio.com) software was used for better quality graphics. The phylogenetic relationship was inferred through MEGA 5.1 (http://www.megasoftware.net/) software. The evolutionary history was inferred using the Neighbor-Joining method, with and percentage of replicate trees in which the associated taxa clustered together in the bootstrap test (1000 replicates). The evolutionary distances were computed using the p-distance method, presented in the units of the number of amino acid differences per site. A work flow for the Phylogenomic analysis has been presented in the supplemental document **S6**. Following major steps were followed:

I. Alignment of orthologous sequences for the selected genes Cysteine Protease, Aquaporin and Alphatubulin using MAFFT v6.864 at default parameters (Auto (FFT-NS-1, FFT-NS-2, FFT-NS-i or L-INS-i) with Amino Acid substitution matrix (BLOSUM62), Gap Penalty (1.53), offset penalty (0.123) and saved in Phylip Interleaved alignment format.
II. Alignment was used to generate RAxML tree, using T-REX online (Boc et al., 2012), at following parameters for generating de-novo phylogeny at following parameters: PROTCATDAYHOFF substitution model, Hill Climbing Alogrithm, Number of alternative runs on distinct starting trees =100, Rapid bootstrap random seed =12345, Bootstrap random seed=12345. This alternate phylogeny was called H1, as compared to commonly accepted Species phylogeny which was called H0 (the null hypothesis).
III. For Delta SSLS estimation, site wise log likelihood values were calculated using (Jobb et al., 2004) for both H0 and H1 phylogeny. Difference in Sitewise Log likelihood was calculated (Delta SSLS= H0-H1), where negative value supports convergent evolution and positive value supports species phylogeny.
IV. For LRT test (Tree Finder), Phylogenetic reconstruction for H0 and H1 was done under WAG substitution model & Likelihood method for identifying best fit protein model with optimized frequencies with Heterogeneity models (G, GI and I). Parametric bootstrapping analysis was done to compare the two evolutionary hypotheses ‘**H0**’ **and** ‘**H1**’. The resulting p-value is the probability that the likelihood ratio simulated under the null hypothesis is less or equal than the observed. Given a level of significance of 5%, a p-value greater than 95% indicates that H1 is better than H0, and a p-value less than 5% indicates that H1 is worse.

**Modeling Procedure & 3D structural prediction analysis:** All structures of representative protein were retrieved from the Protein Data Bank (www.rcsb.org) and aligned using the structure alignment program STAMP (Russell and Barton, 1992). Models using all four structures as template were generated using Modeller9v10 (Sali and Blundell, 1993). 3D representation of the model was prepared in VMD (Visual Molecular dynamics tool) (Humphrey et al., 1996).

### Immunoblot analysis

#### Wheat seedling protein sample Preparation

Wheat seeds were surface sterilized, imbibed for two consecutive days on moist filter pads placed in the glass Petridis, under deprived light, given alternate 16h/8h light/dark cycle for 3 days and then processed as described previously (Close et al., 1989). Briefly, crude protein extract was prepared by homogenization of seeds in Phosphate buffered saline (PBS) buffer with added benzamidine hydrochloride (1mM) and phenylmethylsulfonyl fluoride (PMSF) (1mM) followed by centrifugation at 15,000 rpm for 30 minutes at 4°C. Supernatant was collected to quantify and optimize the protein sample concentration for SDS PAGE with different amount of protein (viz. 20 μg, 50μg, 100μg, 200μg and 400μg). For further experiments 200μg amount was selected as an optimal concentration for Immunoblot analysis.

#### Mosquito developmental stage (Egg, Larva, Pupa) samples

Different stages of mosquito viz. egg, larva, pupa were collected in PBS containing benzamidine hydrochloride (1mM) and phenylmethylsulfonyl fluoride (PMSF) (1mM) protease inhibitors. The collected mosquito whole body samples were homogenized on ice for 10 minutes, followed by centrifugation at 15,000 rpm for 15 min at 4°C. The clean supernatant was collected and quantified for subsequent analysis as described below.

#### Bacterial protein sample

*BL21* cells of E. coli* (2ml) were grown in LB media containing Ampicilin (100 μg/ml) at 37°C till optical density (OD: 600) reached 0.4-0.6. Harvested cells were spinned down at 12000 rpm and re-suspended with 200ul re-suspension buffer containing 50mM NaH_2_PO_4_ pH 8.0, 300mM NaCl, 10mM Imidazole. Cell lysate was then centrifuged at 12000 rpm for 5 min and clear supernatant was analyzed through SDS-PAGE.

### SDS-PAGE and Immunoblot analysis

Protein samples (200μg each) were separated on SDS-polyacrylamide gel with Amersham mini vertical electrophoresis system and transferred to nitrocellulose membrane. Membranes were blocked with 1.5% (w/v) gelatin in PBST and incubated with anti dehydrin primary antibody (1:1000). The unbound antibody was washed three times for 5 min with PBST. Membranes were then incubated with Anti-rabbit HRP secondary antibody (1:60,000) for 1 hour. Unbound secondary antibody was washed for 5 minutes three times with PBST at room temperature. The blots were visualized using Amersham ECL Prime Western blotting detection reagent containing Solution A: luminol enhancer and Solution B: peroxide and developed on X-ray films by developer and bands were readily fixed in fixer solution.

**Genomic DNA isolation & PCR:** For the genomic DNA extraction, a total of five adult female mosquitoes, decapitated with head and wing, were collected in extraction buffer and processed as described earlier (Sharma et al., 2014). All the PCR amplifications conditions and parameters were identical as described above for RT-PCR analysis.

## Conclusion

Evolution and adaptation to dual feeding (sugar vs. blood) behavior of adult female mosquito still remains a central question, a knowledge critical to design vector borne disease management strategies. Comparative salivary transcriptomic and meta-genomics analysis provide initial evidence that mosquito *Anopheles culicifacies*, may have acquired and evolved with plant like machinery components partly shared by salivary associated microbes, together facilitating feeding preference and adaptation over plants grown in the plain agricultural area of rural India.

## Acknowledgement

We are thankful to Dr. S.K. Subbarao for expert comments on the manuscript. We thank Dr. Timothy close, for kind gift of Anti-dehydrin antibody. We thank DBT and ICMR for financial support to conduct the research at NIMR. We thank Kunwarjeet Singh for technical assistance and mosquito rearing.

### Author’s Contribution

Conceived and designed the experiments: RD, PS,NV, KCP. Performed the Experiments: PS, SS, TT,TDD, RD,VK,SLR Analyzed the data: PS, AKM, SS, RD,SV.

Contributed reagents/materials/analysis tools: NS, RD, NV, KCP. Wrote the paper: RD, PS, NV, KCP

### Author Information

The sequence data has been submitted to NCBI SRA database under following accession number: AC-SG-SF: SRR1017392 & AC-SG-BF: SRR1011070. There is no competing financial interest to declare. Correspondence and request for material should be addressed to RD.

## References

Adak, T., Kaur, S. and Singh, O. P. (1999). Comparative susceptibility of different members of the Anopheles culicifacies complex to Plasmodium vivax. Trans R Soc Trop Med Hyg 93, 573–7.

Agrawal, A. A., Hastings, A. P., Johnson, M. T., Maron, J. L. and Salminen, J. P. (2012). Insect herbivores drive real-time ecological and evolutionary change in plant populations. Science 338, 113–6.

Altschul, S. F., Gish, W., Miller, W., Myers, E. W. and Lipman, D. J. (1990). Basic local alignment search tool. J Mol Biol 215, 403–10.

Arendt, J. and Reznick, D. (2008). Convergence and parallelism reconsidered: what have we learned about the genetics of adaptation? Trends Ecol Evol 23, 26–32.

Baldwin, I. T., Halitschke, R., Kessler, A. and Schittko, U. (2001). Merging molecular and ecological approaches in plant-insect interactions. Curr Opin Plant Biol 4, 351–8.

Bao, Y. Y., Wang, Y., Wu, W. J., Zhao, D., Xue, J., Zhang, B. Q., Shen, Z. C. and Zhang, C. X. (2012). De novo intestine-specific transcriptome of the brown planthopper Nilaparvata lugens revealed potential functions in digestion, detoxification and immune response. Genomics 99, 256–64.

Benedict, M. Q. and Seawright, J. A. (1987). Changes in Pigmentation in Mosquitoes (Diptera: Culicidae) in Response to Color of Environment. Annals of the Entomological Society of America 80, 55-61.

Bennett, A. E. (2013). Can plant–microbe–insect interactions enhance or inhibit the spread of invasive species? Functional Ecology 27, 661–671.

Boc, A., Diallo, A. B. and Makarenkov, V. (2012). T-REX: a web server for inferring, validating and visualizing phylogenetic trees and networks. Nucleic Acids Res 40, W573–9.

Borovskii, G. B., Stupnikova, I. V., Antipina, A. I., Vladimirova, S. V. and Voinikov, V. K. (2002). Accumulation of dehydrin-like proteins in the mitochondria of cereals in response to cold, freezing, drought and ABA treatment. BMC Plant Biol 2, 5.

Boto, L. (2014). Horizontal gene transfer in the acquisition of novel traits by metazoans. Proc Biol Sci 281, 20132450.

Campbell, S. A. and Close, T. J. (1997). Dehydrins: genes, proteins, and associations with phenotypic traits. New Phytologist 137, 61–74.

Close, T. J. (1997). Dehydrins: A commonality in the response of plants to dehydration and low temperature. Physiologia Plantarum 100, 291–296.

Close, T. J., Fenton, R. D. and Moonan, F. (1993). A view of plant dehydrins using antibodies specific to the carboxy terminal peptide. Plant Mol Biol 23, 279–86.

Close, T. J., Kortt, A. A. and Chandler, P. M. (1989). A cDNA-based comparison of dehydration-induced proteins (dehydrins) in barley and corn. Plant Mol Biol 13, 95–108.

Close, T. J. and Lammers, P. J. (1993). An osmotic stress protein of cyanobacteria is immunologically related to plant dehydrins. Plant Physiol 101, 773–9.

Conesa, A., Gotz, S., Garcia-Gomez, J. M., Terol, J., Talon, M. and Robles, M. (2005). Blast2GO: a universal tool for annotation, visualization and analysis in functional genomics research. Bioinformatics 21, 3674–6.

Das, S., Radtke, A., Choi, Y. J., Mendes, A. M., Valenzuela, J. G. and Dimopoulos, G. (2010). Transcriptomic and functional analysis of the Anopheles gambiae salivary gland in relation to blood feeding. BMC Genomics 11, 566.

Dixit, R., Rawat, M., Kumar, S., Pandey, K. C., Adak, T. and Sharma, A. (2011). Salivary gland transcriptome analysis in response to sugar feeding in malaria vector Anopheles stephensi. J Insect Physiol 57, 1399–406.

Dixit, R., Sharma, A., Mourya, D. T., Kamaraju, R., Patole, M. S. and Shouche, Y. S. (2009). Salivary gland transcriptome analysis during Plasmodium infection in malaria vector Anopheles stephensi. Int J Infect Dis 13, 636–46.

Ehrlich, P. R. R., P. H. (1964). Butterflies and Plants: A Study in Coevolution. Evolution **Evolution**, 496 586–608.

Eichorst, S. A., Kuske, C. R. and Schmidt, T. M. (2011). Influence of plant polymers on the 498 distribution and cultivation of bacteria in the phylum Acidobacteria. Appl Environ Microbiol 77, 586–96.

Felton, G. W. and Tumlinson, J. H. (2008). Plant-insect dialogs: complex interactions at the 500 plant-insect interface. Curr Opin Plant Biol 11, 457–63.

Ferry, N., Edwards, M. G., Gatehouse, J. A. and Gatehouse, A. M. (2004). Plant-insect 502 interactions: molecular approaches to insect resistance. Curr Opin Biotechnol 15, 155–61.

Foster, W. A. (1995). Mosquito sugar feeding and reproductive energetics. Annu Rev Entomol 40, 443–74.

Fraenkel, G. S. (1959). The raison d’etre of secondary plant substances; these odd chemicals arose as a means of protecting plants from insects and now guide insects to food. Science 129, 1466–70.

Gibbons, J. G., Janson, E. M., Hittinger, C. T., Johnston, M., Abbot, P. and Rokas, A. (2009). Benchmarking next-generation transcriptome sequencing for functional and evolutionary genomics. Mol Biol Evol 26, 2731–44.

Grabherr, M. G. (2013). Trinity: reconstructing a full-length transcriptome without a genome from RNA-Seq data. Nat Biotechnol 29, 644–652.

Grudkowska, M. and Zagdanska, B. (2004). Multifunctional role of plant cysteine proteinases. Acta Biochim Pol 51, 609–24.

Guo, Y., Ribeiro, J. M., Anderson, J. M. and Bour, S. (2009). dCAS: a desktop application for cDNA sequence annotation. Bioinformatics 25, 1195–6.

Hanin, M., Brini, F., Ebel, C., Toda, Y., Takeda, S. and Masmoudi, K. (2011). Plant dehydrins and stress tolerance: versatile proteins for complex mechanisms. Plant Signal Behav 6, 1503–9.

Hittinger, C. T., Johnston, M., Tossberg, J. T. and Rokas, A. (2010). Leveraging skewed transcript abundance by RNA-Seq to increase the genomic depth of the tree of life. Proc Natl Acad Sci U S A 107, 520 1476–81.

Hogenhout, S. A. and Bos, J. I. (2011). Effector proteins that modulate plant--insect interactions. Curr Opin Plant Biol 14, 422–8.

Humphrey, W., Dalke, A. and Schulten, K. (1996). VMD: visual molecular dynamics. J Mol Graph 14, 33–8, 27–8.

James, A. A. (2003). Blocking malaria parasite invasion of mosquito salivary glands. J Exp Biol 206, 3817–21.

Jobb, G., von Haeseler, A. and Strimmer, K. (2004). TREEFINDER: a powerful graphical analysis environment for molecular phylogenetics. BMC Evol Biol 4, 18.

Juhn, J., Naeem-Ullah, U., Maciel Guedes, B. A., Majid, A., Coleman, J., Paolucci Pimenta, P. F., Akram, W., James, A. A. and Marinotti, O. (2011). Spatial mapping of gene expression in the salivary glands of the dengue vector mosquito, Aedes aegypti. Parasit Vectors 4, 1.

Larkin, M. A., Blackshields, G., Brown, N. P., Chenna, R., McGettigan, P. A., McWilliam, H., Valentin, F., Wallace, I. M., Wilm, A., Lopez, R. et al. (2007). Clustal W and Clustal X version 2.0. Bioinformatics 23, 2947–8.

Lehane, J. (2005). The Biology of Blood-Sucking in Insects: Cambridge University Press.

Li, R., Brawley, S. H. and Close, T. J. (1998). PROTEINS IMMUNOLOGICALLY RELATED TO DEHYDRINS IN FUCOID ALGAE. Journal of Phycology 34, 642–650.

Livak, K. J. and Schmittgen, T. D. (2001). Analysis of relative gene expression data using real-time quantitative PCR and the 2(-Delta Delta C(T)) Method. Methods 25, 402–8.

Lu, Y., Park, Y., Gao, X., Zhang, X., Yao, J., Pang, Y. P., Jiang, H. and Zhu, K. Y. (2012). Cholinergic and non-cholinergic functions of two acetylcholinesterase genes revealed by gene-silencing in Tribolium castaneum. Sci Rep 2, 288.

Maffei, M. E., Mithofer, A. and Boland, W. (2007). Before gene expression: early events in plant–insect interaction. Trends Plant Sci 12, 310–6.

Maurel, C., Verdoucq, L., Luu, D. T. and Santoni, V. (2008). Plant aquaporins: membrane channels with multiple integrated functions. Annu Rev Plant Biol 59, 595–624.

McQueen-Mason, S. J. and Cosgrove, D. J. (1995). Expansin mode of action on cell walls. Analysis of wall hydrolysis, stress relaxation, and binding. Plant Physiol 107, 87–100.

Moran, N. A. and Jarvik, T. (2010). Lateral transfer of genes from fungi underlies carotenoid production in aphids. Science 328, 624–7.

Mtwisha, L., Brandt, W., McCready, S. and Lindsey, G. G. (1998). HSP 12 is a LEA-like protein in Saccharomyces cerevisiae. Plant Mol Biol 37, 513–21.

Neafsey, D. E., Christophides, G. K., Collins, F. H., Emrich, S. J., Fontaine, M. C., Gelbart, W., Hahn, M. W., Howell, P. I., Kafatos, F. C., Lawson, D. et al. (2013). The evolution of the Anopheles 16 genomes project. G3 (Bethesda) 3, 1191–4.

Ouellet, F., Houde, M. and Sarhan, F. (1993). Purification, characterization and cDNA cloning of the 200 kDa protein induced by cold acclimation in wheat. Plant Cell Physiol 34, 59–65.

Parker, J., Tsagkogeorga, G., Cotton, J. A., Liu, Y., Provero, P., Stupka, E. and Rossiter, S. J. (2013). Genome-wide signatures of convergent evolution in echolocating mammals. Nature 502, 228–31.

Pieterse, C. M. and Dicke, M. (2007). Plant interactions with microbes and insects: from molecular mechanisms to ecology. Trends Plant Sci 12, 564–9.

Rai, K. S. (1999). Genetics of mosquitoes. Journal of Genetics 78, 163–169.

Reid, N. M., Addison, S. L., Macdonald, L. J. and Lloyd-Jones, G. (2011). Biodiversity of active and inactive bacteria in the gut flora of wood-feeding huhu beetle larvae (Prionoplus reticularis). Appl Environ Microbiol 77, 7000–6.

Ribeiro, J. M., Mans, B. J. and Arca, B. (2010). An insight into the sialome of blood-feeding Nematocera. Insect Biochem Mol Biol 40, 767–84.

Rodrigues, J., Brayner, F. A., Alves, L. C., Dixit, R. and Barillas-Mury, C. (2010). Hemocyte differentiation mediates innate immune memory in Anopheles gambiae mosquitoes. Science 329, 1353–5.

Rodriguez, M. H. and Hernandez-Hernandez Fde, L. (2004). Insect-malaria parasites interactions: the salivary gland. Insect Biochem Mol Biol 34, 615–24.

Russell, R. B. and Barton, G. J. (1992). Multiple protein sequence alignment from tertiary structure comparison: assignment of global and residue confidence levels. Proteins 14, 309–23.

Sali, A. and Blundell, T. L. (1993). Comparative protein modelling by satisfaction of spatial restraints. J Mol Biol 234, 779–815.

Scott, J. G., Michel, K., Bartholomay, L. C., Siegfried, B. D., Hunter, W. B., Smagghe, G., Zhu, K. Y. and Douglas, A. E. (2013). Towards the elements of successful insect RNAi. J Insect Physiol 59, 1212–21.

Sharma, P., Sharma, S., Maurya, R. K., De, T. D., Thomas, T., Lata, S., Singh, N., Pandey, K. C., Valecha, N. and Dixit, R. (2014). Salivary glands harbor more diverse microbial communities than gut in Anopheles culicifacies. Parasit Vectors 7, 235.

Su, Y. L., Li, J. M., Li, M., Luan, J. B., Ye, X. D., Wang, X. W. and Liu, S. S. (2012). Transcriptomic analysis of the salivary glands of an invasive whitefly. PLoS One 7, e39303.

Takahashi, R., Joshee, N. and Kitagawa, Y. (1994). Induction of chilling resistance by water stress, and cDNA sequence analysis and expression of water stress-regulated genes in rice. Plant Mol Biol 26, 339–52.

Valmalette, J. C., Dombrovsky, A., Brat, P., Mertz, C., Capovilla, M. and Robichon, A. (2012). Light-induced electron transfer and ATP synthesis in a carotene synthesizing insect. Sci Rep 2, 579.

Venancio, T. M., Cristofoletti, P. T., Ferreira, C., Verjovski-Almeida, S. and Terra, W. R. (2009). The Aedes aegypti larval transcriptome: a comparative perspective with emphasis on trypsins and the domain structure of peritrophins. Insect Mol Biol 18, 33–44.

Wang, X. W., Luan, J. B., Li, J. M., Bao, Y. Y., Zhang, C. X. and Liu, S. S. (2010). De novo characterization of a whitefly transcriptome and analysis of its gene expression during development. BMC Genomics 11, 400.

Xie, C., Mao, X., Huang, J., Ding, Y., Wu, J., Dong, S., Kong, L., Gao, G., Li, C. Y. and Wei, L. (2011). KOBAS 2.0: a web server for annotation and identification of enriched pathways and diseases. Nucleic Acids Res 39, W316–22.

Zhen, Y., Aardema, M. L., Medina, E. M., Schumer, M. and Andolfatto, P. (2012). Parallel molecular evolution in an herbivore community. Science 337, 1634–7.

